# Characterization of various remdesivir-resistant mutations of SARS-CoV-2 by mathematical modeling and molecular dynamics simulation

**DOI:** 10.1101/2022.02.22.481436

**Authors:** Shiho Torii, Kwang Su Kim, Jun Koseki, Rigel Suzuki, Shoya Iwanami, Yasuhisa Fujita, Yong Dam Jeong, Yoshiharu Matsuura, Teppei Shimamura, Shingo Iwami, Takasuke Fukuhara

## Abstract

Mutations continue to accumulate within the SARS-CoV-2 genome, and the ongoing epidemic has shown no signs of ending. It is critical to predict problematic mutations that may arise in clinical environments and assess their properties in advance to quickly implement countermeasures against future variant infections. In this study, we identified mutations resistant to remdesivir, which is widely administered to SARS-CoV-2-infected patients, and discuss the cause of resistance. First, we simultaneously constructed eight recombinant viruses carrying the mutations detected in *in vitro* serial passages of SARS-CoV-2 in the presence of remdesivir. Time course analyses of cellular virus infections showed significantly higher infectious titers and infection rates in mutant viruses than wild type virus under treatment with remdesivir. Next, we developed a mathematical model in consideration of the changing dynamic of cells infected with mutant viruses with distinct propagation properties and defined that mutations detected in *in vitro* passages canceled the antiviral activities of remdesivir without raising virus production capacity. Finally, molecular dynamics simulations of the NSP12 protein of SARS-CoV-2 revealed that the molecular vibration around the RNA-binding site was increased by the introduction of mutations on NSP12. Taken together, we identified multiple mutations that affected the flexibility of the RNA binding site and decreased the antiviral activity of remdesivir. Our new insights will contribute to developing further antiviral measures against SARS-CoV-2 infection.

**Significance Statement:** Considering the emerging Omicron strain, quick characterization of SARS-CoV-2 mutations is important. However, owing to the difficulties in genetically modifying SARS-CoV-2, limited groups have produced multiple mutant viruses. Our cutting-edge reverse genetics technique enabled construction of eight reporter-carrying mutant SARS-CoV-2 in this study. We developed a mathematical model taking into account sequential changes and identified antiviral effects against mutant viruses with differing propagation capacities and lethal effects on cells. In addition to identifying the positions of mutations, we analyzed the structural changes in SARS-CoV-2 NSP12 by computer simulation to understand the mechanism of resistance. This multidisciplinary approach promotes the evaluation of future resistance mutations.

## Introduction

Severe acute respiratory syndrome coronavirus 2 (SARS-CoV-2) was first discovered in 2019 and quickly spread around the world (*1*). Novel SARS-CoV-2 variants have since continued to emerge and the number of virus-infected cases repeats increases and decreases (*2*). The clinical spectrum of SARS-CoV-2 infection ranges from mild to critical. While most infections present mild or minor symptoms (e.g. fever, cough, sore throat, malaise, headache, muscle pain, nausea, vomiting, diarrhea, loss of taste and smell), severe acute respiratory disease requires admission to intensive care (*3–5*). The illness can be observed even after successful vaccination (*6*). Antiviral drugs that can be administered to patients after moderate or severe clinical symptoms have been observed have played important roles in clinical environments. Therefore, it is vital to understand the effectiveness of currently approved antivirals from multiple angles to develop future drugs. In particular, the potential to drive drug resistance should be evaluated because drug-resistant mutations have been observed in several viruses such as influenza A virus, human immunodeficiency virus and hepatitis B virus in the clinical environment (*7–10*).

Remdesivir (RDV) (GS-5734) is the US Food and Drug Administration (FDA)-approved drug for treatment of coronavirus disease 2019 (COVID-19) patients (*11, 12*). The compound is an intravenously administered adenosine analogue prodrug that binds to the viral RNA-dependent RNA polymerase and inhibits viral replication. It has demonstrated antiviral activities against a broad range of RNA viruses including Ebolavirus, SARS-CoV, MERS-CoV and SARS-CoV-2 (*13–17*). RDV has been widely used in the treatment of SARS-CoV-2 patients, however only two amino acid mutations (D484Y and E802D in non-structural protein [NSP]12) were identified from SARS-CoV-2 patients that were administered RDV (*18, 19*). One mutation (E802D) was also found in *in vitro* serial passages of the virus under treatment of RDV (*20*). Although studies regarding E802D revealed that the mutation decreased viral susceptibility to RDV (*19, 20*), the mechanisms of how resistance arises have not yet been analyzed in detail. It is critical to elucidate the mechanisms of RDV resistance and to identify further RDV-resistant mutants that may arise in the future to circumvent resistance mutations before they become established in circulating strains.

To evaluate the effect of each gene mutation on viral propagation, genetically modified viruses should be engineered using the reverse genetics system. We recently established a quick reverse genetics system for SARS-CoV-2 using the circular polymerase extension reaction (CPER) method (*21*). Nine viral genome fragments, which cover the full-length viral genome, and a linker fragment that encodes the promoter sequence were amplified by PCR and connected to obtain the circular viral DNAs by an additional PCR. By direct transfection of the circular DNAs, infectious SARS-CoV- 2 was rescued. Introduction of reporters or mutations can be quickly completed by overlapping PCR or plasmid mutagenesis using the desired gene fragments of less than 5,000 base pairs (bp). While other reverse genetics systems for SARS-CoV-2 require specific techniques such as *in vitro* transcription or *in vitro* ligation, which are obstacles to mutagenesis (*22, 23*), our method does not need these and has already been applied to the characterization of several viral mutations observed in the different SARS-CoV-2 variants (*24, 25*), allowing us to simultaneously generate multiple mutants (*26*).

In this study, we attempted to identify multiple RDV-resistant mutations and examine the mechanisms of RDV resistance by a multidisciplinary approach that integrates state-of-the-art reverse genetics, mathematical modeling, and molecular dynamics analyses. We first predicted the presumed RDV-resistant mutations by *in vitro* passages of SARS-CoV-2 in the presence of RDV. Next, the recombinant viruses carrying the predicted mutations were generated by the CPER method and the efficiency of infectious virus production and antiviral effects of RDV on the mutants were examined by mathematical modeling. Finally, the conformational changes of NSP12 induced by mutations were analyzed by molecular dynamics simulations to understand the mechanisms of RDV resistance.

## Results

### Generation of RDV-resistant SARS-CoV-2

To identify the genes presumably involved in RDV resistance, we first passaged the SARS-CoV-2 strain JPN/TY/WK-521 in HEK293-C34 cells under treatment with RDV (Fig. 1A). The concentration of RDV was 0.01 μM in the first passage (P1) and was gradually increased over 10 passages. Throughout the passages, the virus-infected cells were cultured until cytopathic effect (CPE) was observed (3–8 days). No CPE was observed during 14 days treatment with 0.1 μM RDV in P1, indicating that the drug was effective at suppressing infection at this low concentration. However, CPE was observed throughout the wells in the presence of 4.0 μM RDV at P10, indicating that the virus decreased susceptibility to RDV during the passages. After 10 passages, the culture supernatants were collected and subjected to Sanger sequencing to determine the full-length viral sequence. Comparison of the P10 virus sequence with the original SARS-CoV-2 genome found six unique mutation sites (Fig. 1B). The deletion of nine nucleotides was observed in NSP1, and amino acid substitutions were observed in NSP4, NSP6, NSP12 and NSP15. According to Nextstrain (*27*), the same mutations had been detected in NSP1, NSP4, NSP6 and NSP15, but only a few cases of each mutation had been reported. To date, there have been no reports of the E796G or C799F mutations in NSP12 identified here.

**Fig. 1.**
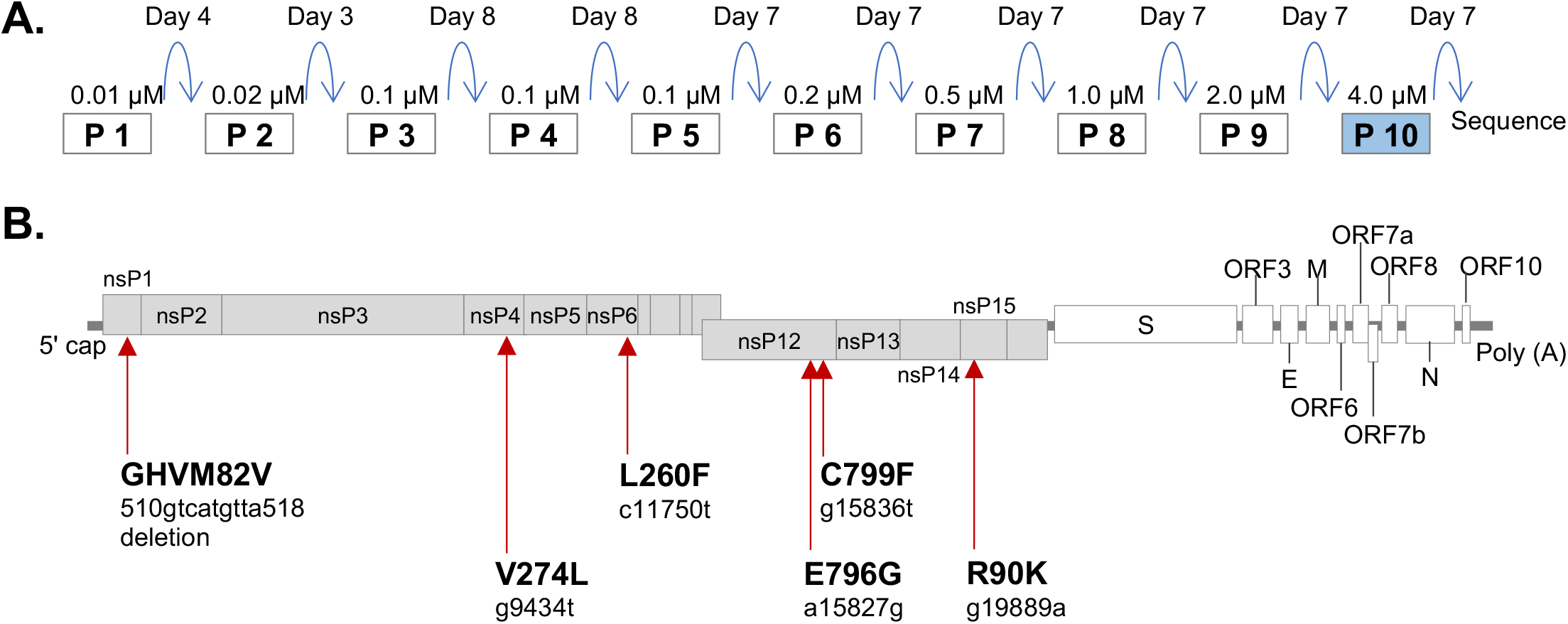
Identification of RDV-resistant mutations of SARS-CoV-2. **(A)** Schematic image of the *in vitro* serial passage of SARS-CoV-2 in the presence of RDV. Supernatants of virus-infected cells were passaged with gradually increased concentrations of RDV. The virus sequence was examined after 10 passages. **(B)** The SARS-CoV-2 genome with the locations of mutations observed in the in vitro serial passages.

We then generated high-affinity NanoBiT (HiBiT)-carrying recombinant SARS-CoV-2 with each mutation to identify the RDV-resistant mutations. NanoLuc enzymatic activity can be detected by interaction of HiBiT and large NanoBiT (LgBiT), which constitute a split reporter. The reporter SARS-CoV-2 can be generated by inserting only 11 amino acids into the viral genome, and HiBiT- carrying viruses exhibit similar growth kinetics to wildtype (WT) virus (*21*). All recombinant SARS-CoV-2 with HiBiT and mutations were prepared using the CPER method that was previously established by our group. Amino acid substitutions were introduced by overlapping PCR and the full-length sequences of the mutant viruses were confirmed prior to assay by Sanger sequencing.

Because RDV acts as a nucleoside analog and targets the RNA-dependent RNA polymerase (RdRp) of coronaviruses, including SARS-CoV-2, in the current study we focused on the mutations in NSP12. We generated recombinant viruses with the E796G or C799F mutations that were observed in our P10 serial virus passage (Table 1). We also prepared recombinant viruses with all mutations observed after P10 (R10/C799F/E796G) or with all mutations except for E796G (R10/C799F). In addition to the mutations observed in this study, we also characterized mutations that have been reported as, or anticipated to be, resistant to RDV, as listed in Table 1.

**Table 1.**
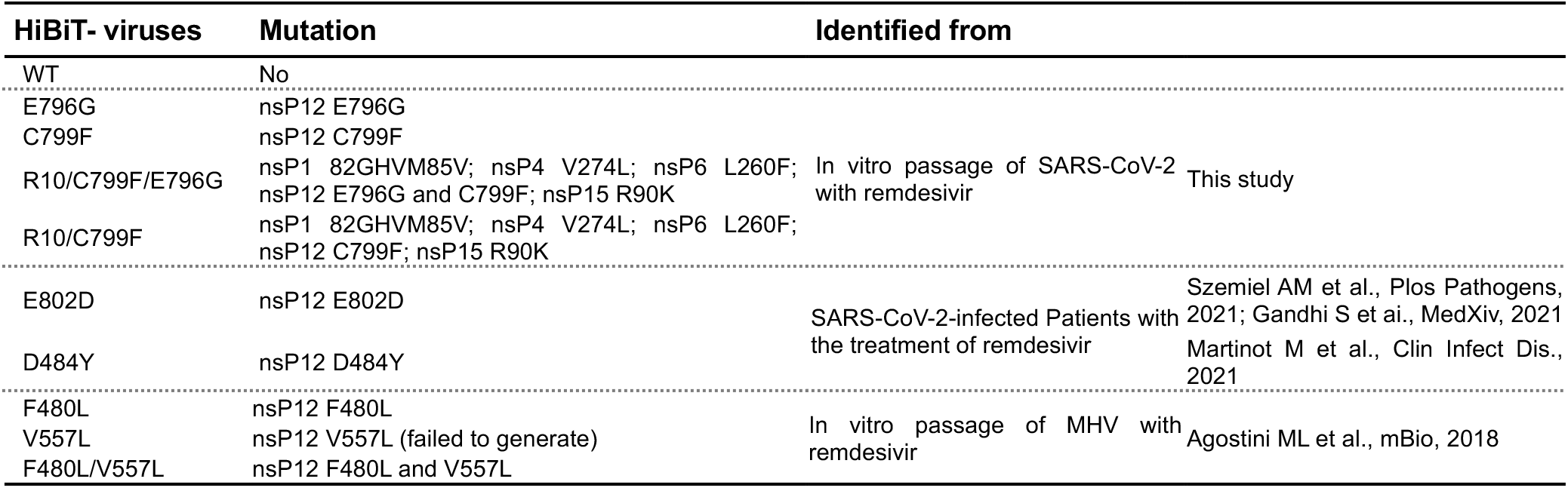
Generation of the recombinant SARS-CoV-2 carrying HiBiT reporter and mutations

The amino acid mutation E802D in NSP12 was found during the serial passage of SARS-CoV- 2 *in vitro* in the presence of RDV and another report showed that the same mutation was found in patients receiving RDV (*19, 20*). The D484Y mutation was also identified in a COVID-19 patient receiving RDV treatment (*18*). Previously, amino acid substitutions F476L and V553L were identified as RDV-resistant mutations in the *Betacoronavirus* murine hepatitis virus (MHV)(*15*). The two affected amino acid residues (476F and 553L in MHV) are conserved across coronaviruses and correspond to 480F and 557V in the SARS-CoV-2 genome. We attempted to generate recombinant SARS-CoV-2 with either or both mutations but the virus with the single V557L mutation could not be rescued.

We then examined the sensitivity of recombinant viruses to RDV (Fig. S1). Viruses were cultured in the presence of RDV at 0–1.0 μM final concentration for 48 hours and luciferase activity was measured and normalized against control without RDV treatment (0 μM final concentration). The 50% effective concentration (EC50) was calculated using the drc package (v3.0-1; R Project for Statistical Computing). All tested mutant viruses showed greater EC50 than WT virus, although the difference between WT and F480L mutant was small, indicating that the mutations observed in NSP12 led to decreased viral sensitivity to RDV.

### Time course analyses of infection with presumed RDV-resistant mutants

To characterize the growth efficiency of mutant viruses and the antiviral effects of RDV, we first performed time course analyses of infectious virus production with or without RDV for 72 hours (Fig. 2A and B). At all the indicated time points, the infectious titer of each mutant virus was similar or lower than that of the WT virus in the absence of RDV treatment, indicating that the mutant viruses produced the infectious viruses with the same or lower efficiency as WT virus (Fig. 2A). Conversely, significant differences were observed in the infectious titers of mutant viruses in the presence of RDV at 0.05 μM final concentration. In the left panel of Fig. 2B, the infectious titers of mutant viruses (E796G, C799F, R10/E796G/C799F, and R10/C799F) gradually increased for 48 hours and were significantly higher than that of WT virus at 72 hours post infection (hpi). In the right panel of Fig. 2B, the infectious titer of the E802D mutant virus increased rapidly and was significantly higher than that of WT virus at 48 and 72 hpi. The titers of the F484Y and F480L/V557L mutants were also significantly higher than that of WT virus. Meanwhile there were no differences between the titers of F480L mutant and WT viruses at the indicated time points. These results suggest that the sensitivity of all the mutant viruses, except for the F480L virus, to RDV was diminished, which was consistent with the results of the RDV susceptibility test demonstrating minimal change in the EC50 of the F480L virus (Fig. S1).

**Fig. 2.**
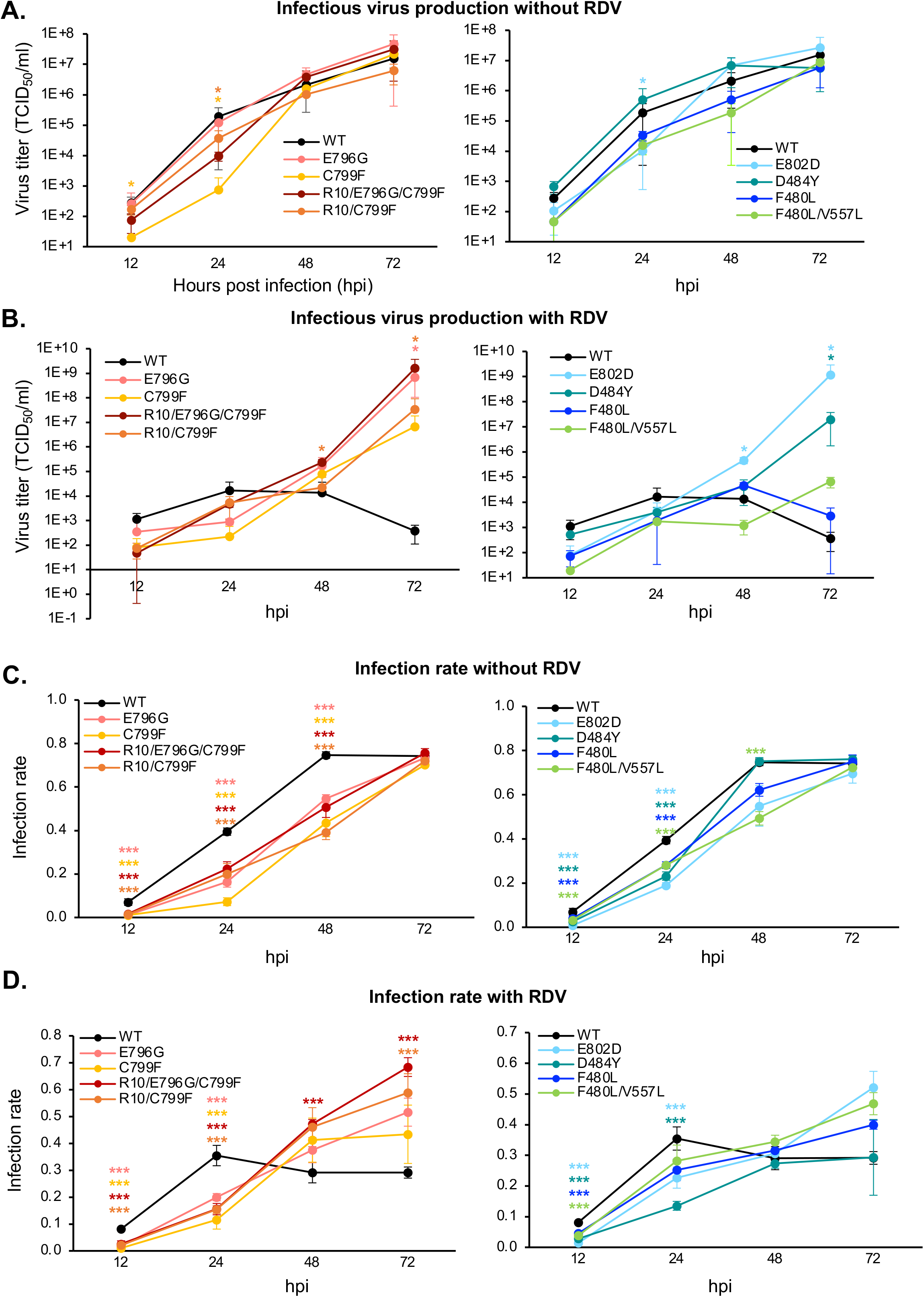
Infection kinetics of recombinant SARS-CoV-2 with NSP12 mutations. (A and B) Infectious virus production in the absence **(A)** and presence **(B)** of RDV. Supernatants of virus- infected cells were collected from 12–96 hpi and virus titers were determined by titration. Statistical significances were determined by Kruskal–Wallis test with the two stage linear step-up procedure of Benjamini, Kreiger and Yekutieli. Significant differences compared with WT virus are indicated by an asterisk (\**p<0.05*). **(C and D)** Infection rate in the absence **(C)** and presence **(D)** of RDV. Virus-infected cells were fixed and stained with antibodies for SARS-CoV-2 and cell nuclei. Infection rates were calculated by dividing the numbers of virus-positive cells by the numbers of nuclei. Statistical significances were determined by one-way ANOVA with Dunnett’s test. Significant differences compared with WT virus are indicated by asterisks (\*\*\**p<0.001*).

Next, we investigated the the ratio of the virus-infected cells (Fig. 2C and D). HEK293-C34 cells were infected with mutant viruses with and without RDV treatment. Virus-infected cells were harvested and fixed from 12–72 hpi and subjected to immunofluorescent assay using anti-SARS-CoV-2 NP antibody and DAPI. The virus infection rates were then calculated. All the mutant viruses demonstrated equivalent or significantly lower virus infection rates compared with WT virus in the absence of RDV. These data suggested that the number of cells infected with mutant viruses increased more slowly compared with WT virus in the absence of RDV treatment, consistent with the data on production of infectious virus particles. Meanwhile, the infection rates of the presumed RDV-resistant mutant viruses, except for D484Y, were higher or significantly higher (R10/E796G/C799F at 48 and 72 hpi, and R10/C799F at 72 hpi) than those of WT virus at 48 and 72 hpi in the presence of RDV, indicating that these mutant viruses can spread more efficiently than WT virus in the presence of RDV.

### Antiviral effect of RDV on the presumed RDV-resistant mutants, analyzed by mathematical modeling

To quantify the kinetic parameters of SARS-CoV-2 and the antiviral effect of RDV on WT and RDV-resistant viruses, we developed a mathematical model for SARS-CoV-2 infection under RDV treatment. We examined the growth rate of HEK293-C34 cells up to 48 hours after seeding (Fig. S2A), the degradation rate of SARS-CoV-2 at 37°C (Fig. S2B), the infectious virus production rate for 96 hours (Fig. 2A and B), and the rate of infection in susceptible cells (Fig. 2C and D). These estimated parameters were fixed and used here.

To consider the variability of kinetic parameters and model predictions, we performed Bayesian estimation for the whole dataset using Markov chain Monte Carlo (MCMC) sampling, and simultaneously fit equations (1–4) with RDV (*ε* > **0**) and without RDV (*ε* = **0**) to the concentrations of target cells, infected cells, and infectious virus (see **Method** and **Fig. S3**). The estimated parameters are listed in **Table 2** and **Table 3**, and the simulation results of the model using these best-fit parameter estimates are shown with the data in **Fig S3**. Comparing the virus production rate and the antiviral effect for the eight different resistant mutants (D484Y, F480L, F480L/V557L, E796G, C799F, R10/C799F, R10/E796G/C799F, E802D), we found that the virus production for all mutants was lower than that of the WT virus except for D484Y mutant (**Fig 3B** and **Table 3**). The D484Y mutation had a competitive advantage in virus production rate, and this property might be involved in its decreased susceptibility to RDV, although the difference from WT was only 1–1.25- fold.

**Fig. 3.**
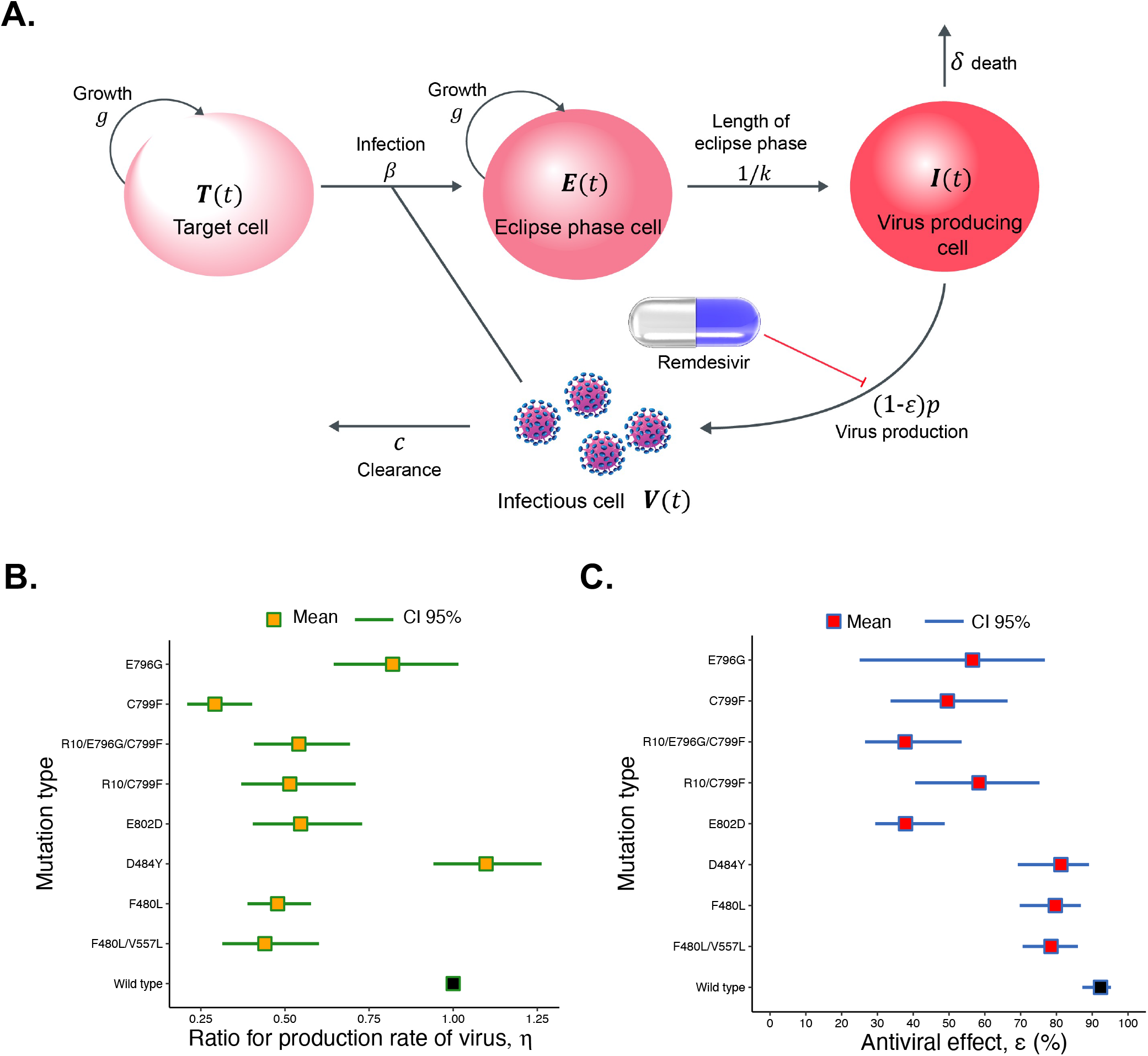
Quantifying antiviral effects of RDV against SARS-CoV-2 infection. **(A)** Schematic diagram of SARS-CoV-2 infection under RDV treatment in cell culture. The target cells are infected by infectious virus at rate ***β*** and become eclipse phase cells. The average duration of the eclipse phase is **1**/***k*** days and these eclipse phase cells start producing viruses at rate ***p*** (i.e., become virus-producing cells). The target cells and eclipse phase cells are assumed to divide at rate ***g***, and virus-producing cells die at rate 𝜹. Progeny infectious viruses are cleared at rate 𝒄. RDV blocks virus production by inhibiting viral replication in infected cells with inhibition rate *ε*. **(B)** Comparison of the fold-change of virus production rate, 𝜼, for eight different RDV-resistant viruses and WT virus. Orange dots are the mean value and green lines show 95% CI, which were estimated from the experiments with/without RDV. For WT, we consider 𝜼 = **1**, shown by the black dot. **(C)** Comparison of the antiviral effect of RDV, *ε*, for eight different RDV-resistant viruses and WT virus.Red and black dots are the mean values of the antiviral effects and blue lines show 95% CI, which were estimated from infection experiments with/without RDV.

**Table 2.**
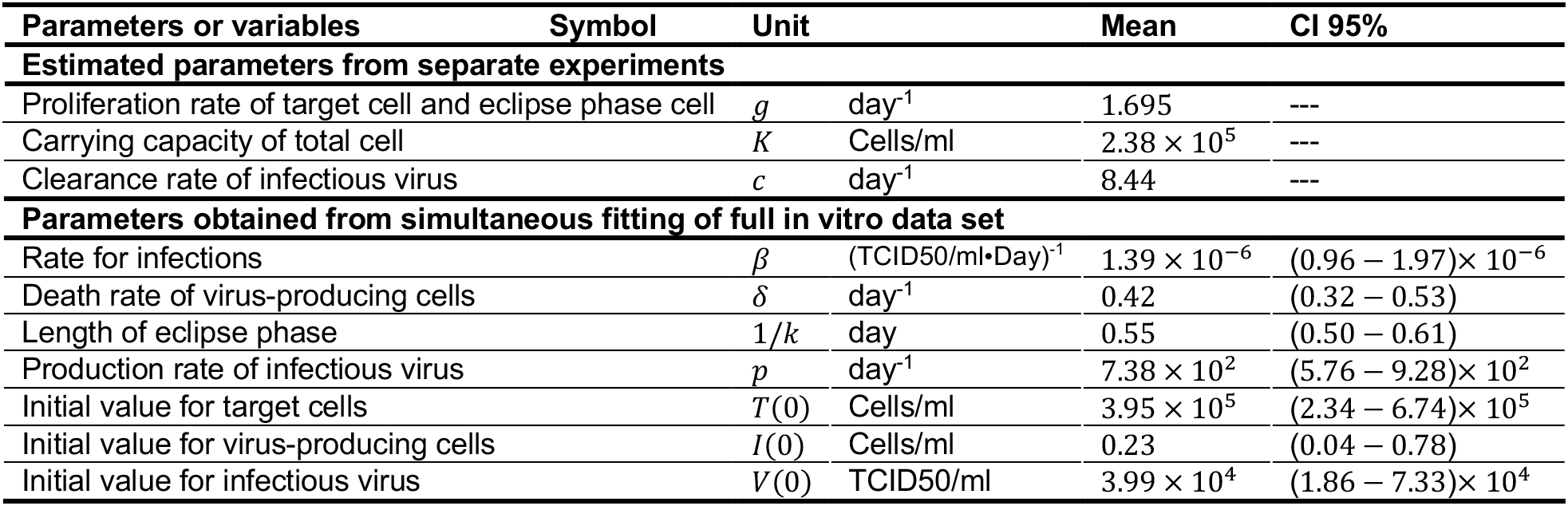
Estimated kinetic parameters and initial values

| Parameters or variables | Symbol | Unit | Mean | CI 95% |
| --- | --- | --- | --- | --- |
| **Estimated parameters from separate experiments** |  |  |  |  |
| Proliferation rate of target cell and eclipse phase cell | $g$ | day<sup>-1</sup> | 1.695 | --- |
| Carrying capacity of total cell | $K$ | Cells/ml | $2.38 \times 10^5$ | --- |
| Clearance rate of infectious virus | $c$ | day<sup>-1</sup> | 8.44 | --- |
| **Parameters obtained from simultaneous fitting of full in vitro data set** |  |  |  |  |
| Rate for infections | $\beta$ | (TCID50/ml·Day)<sup>-1</sup> | $1.39 \times 10^{-6}$ | $(0.96 - 1.97) \times 10^{-6}$ |
| Death rate of virus-producing cells | $\delta$ | day<sup>-1</sup> | 0.42 | (0.32 – 0.53) |
| Length of eclipse phase | $1/k$ | day | 0.55 | (0.50 – 0.61) |
| Production rate of infectious virus | $p$ | day<sup>-1</sup> | $7.38 \times 10^2$ | $(5.76 - 9.28) \times 10^2$ |
| Initial value for target cells | $T(0)$ | Cells/ml | $3.95 \times 10^5$ | $(2.34 - 6.74) \times 10^5$ |
| Initial value for virus-producing cells | $I(0)$ | Cells/ml | 0.23 | (0.04 – 0.78) |
| Initial value for infectious virus | $V(0)$ | TCID50/ml | $3.99 \times 10^4$ | $(1.86 - 7.33) \times 10^4$ |

**Table 3.** Estimated parameters of the virus production and the antiviral effect for eight different resistant mutants and wild type

| Parameters or variables (symbol) | Wild type | E796G | C799F | R10/E796G/C799F | R10/C799F | E802D | D484Y | F480L | F480L/V557L |
| --- | --- | --- | --- | --- | --- | --- | --- | --- | --- |
| Fold-change of the production rate of infectious virus ($\eta$) | 1<br>(---) | 0.82<br>(0.64 – 1.02) | 0.29<br>(0.21 – 0.40) | 0.54<br>(0.41 – 0.69) | 0.51<br>(0.37 – 0.71) | 0.55<br>(0.40 – 0.73) | 1.10<br>(0.94 – 1.26) | 0.48<br>(0.39 – 0.58) | 0.44<br>(0.31 – 0.60) |
| Inhibition rate of virus production by remdesivir ($\varepsilon$) | 0.92<sup>†</sup><br>(0.87 – 0.95)<sup>‡</sup> | 0.57<br>(0.25 – 0.77) | 0.50<br>(0.34 – 0.66) | 0.38<br>(0.27 – 0.54) | 0.58<br>(0.41 – 0.75) | 0.38<br>(0.29 – 0.49) | 0.81<br>(0.69 – 0.89) | 0.80<br>(0.70 – 0.87) | 0.78<br>(0.71 – 0.86) |

<sup>†</sup> Mean value <sup>‡</sup> 95% confidence interval

Interestingly, all the other tested mutations strongly suppressed virus production. RDV showed more than 70% antiviral effect on three mutations, 85% (95% CI: 80–90) for D484Y; 82% (95% CI: 76–89) for F480L; and 76% (95% CI: 68–82) for F480L/V557L. However, the antiviral effect on two mutations was less than 30%, at 28% (95% CI: 22–34) for E802D and 27% (95% CI: 22–34) for R10/E796G/C799F (**Fig 3C** and **Table 3**). All the examined mutations reduced the antiviral effect of RDV and the change was greatest in the mutations found in *in vitro* passages. The antiviral effect of RDV in the F480L mutation was not much different from that observed with WT virus, which is consistent with the results of the RDV susceptibility test. Because there was no correlation between the efficiency of virus production and the antiviral effect, the mechanisms of RDV resistance were predicted to be a structural defect in the direct interaction between the viral genome replication complex and RDV.

### RDV-resistance mechanisms of the presumed RDV-resistant mutants, analyzed by computer simulation

Molecular modeling and molecular dynamics simulations were performed to clarify the structural and property changes caused by amino acid mutations in the NSP12 protein. The representative complete structure of the prepared protein-RNA complex is shown in Fig. 4A. In this figure, NSP12, binding RNA, and RDV are shown in cartoon, ball and stick, and van der Waals notation, respectively. RDV is located at the end of the binding RNA and is inside the protein. Then, the root mean square deviations (RMSDs) were compared using molecular dynamics simulation trajectories of each complex of WT or mutant NSP12 protein and RNA-incorporated RDV, as shown in Fig. 4B. Most complexes reached thermodynamic stability and plateau in RMSD after 200,000 steps. However, only the V557L mutant structure failed to reach the stabilized structure. When RNA structures for this mutant were superimposed and RMSDs were calculated for RNA and protein separately, the increase in RMSDs for RNA reached a plateau, while the RMSDs for protein continued to increase as before. This behavior means the bonds between protein and RNA tended to move apart. In other words, the RNA-protein complex tended to be unstable, which may correspond with the inability of this mutant virus to multiply, indicating the probable reason for failed rescue of the V557L mutant recombinant SARS-CoV-2 (Table 1). Therefore, we proceeded with the analysis of the other mutants only.

**Fig. 4.**
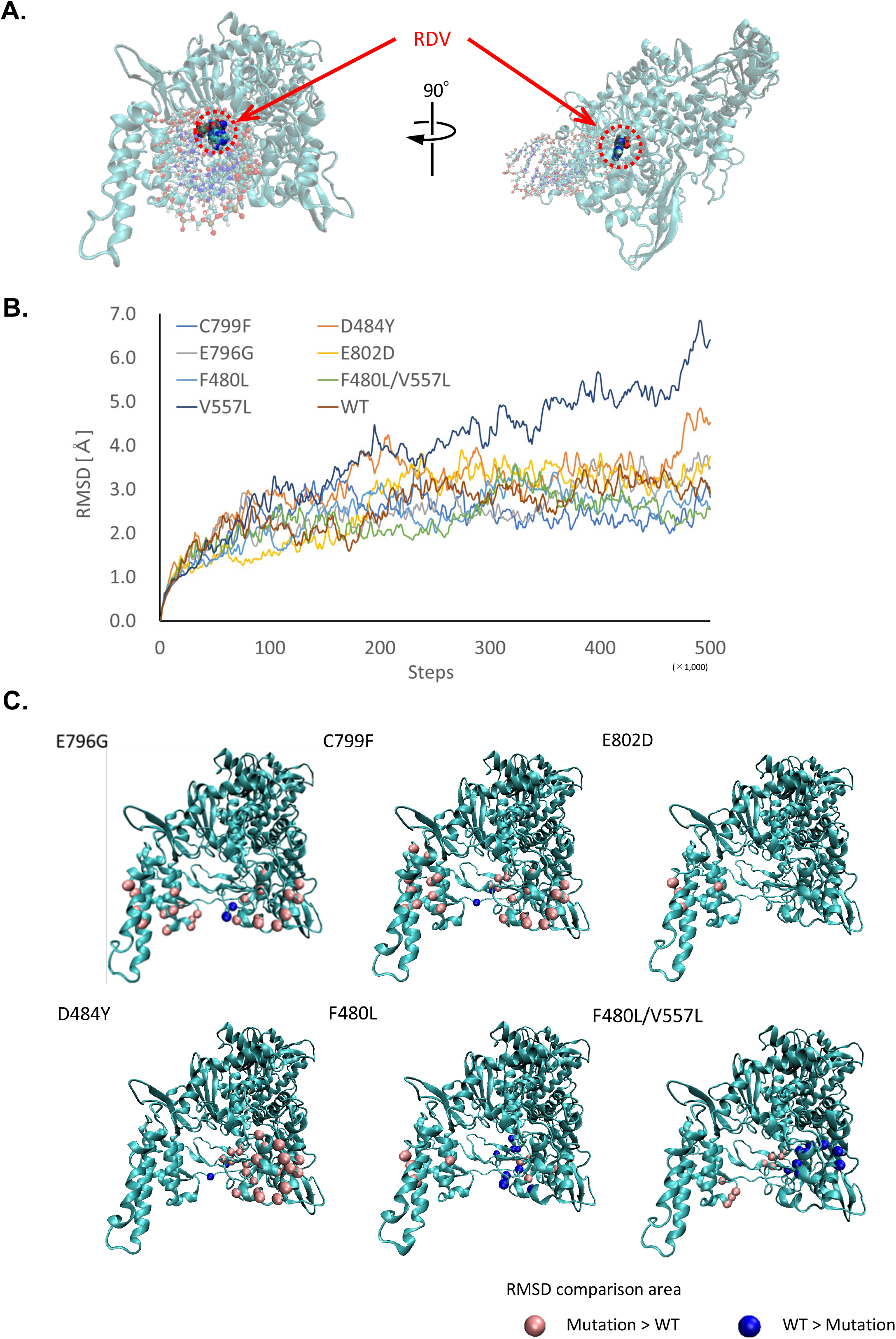
NSP12-RNA binding structure and comparison of thermodynamic stability. **(A)** Overall view of NSP12 protein and location of the RDV. **(B)** RMSD comparison of RNA-binding proteins. **(C)** Comparison of the molecular vibrations of WT and each mutant. The pink and blue spheres represent regions of large oscillations in the mutant and WT, respectively.

Using the trajectories obtained from molecular dynamics calculations, we calculated and compared the variation of RMSD for each substructure. The RMSDs of each mutant complex and the WT complex were searched for areas where they differed significantly, and the results are shown in Fig. 4C. Locations where the RMSD variation of the mutant is greater than that of the WT are indicated by pink spheres, and conversely, locations where the RMSD variation of the WT is greater than that of the mutant are indicated by blue spheres. In the tested mutations, except for F480L, the molecular vibration of the mutants tended to increase around the RNA-binding site as shown in Fig. S4, indicating that introduction of the mutations increased the flexibility of the RNA binding site. However, in the center of the RNA-binding site (near the RDV-binding site) of the F480L mutant, the molecular vibration of the mutant tended to be small, which is consistent with the small change in antiviral effect (Fig. 3C) and RDV sensitivity (Fig. S1).

Taken together, NSP12 mutations found in previous studies and in our *in vitro* virus passages decreased the antiviral effect of RDV, though not to the same degree, and influenced increased flexibility of the RNA-binding site of the NSP12 structure.

## Discussion

As demonstrated by the recent emergence of the Omicron strain, mutations have continuously accumulated on the SARS-CoV-2 genome (*28*). To implement the most effective measures against COVID-19, it is crucial to predict clinically important mutations in advance. In this study, we generated eight recombinant viruses with presumed RDV-resistant mutations, using a simple reverse genetics system. Quantifying the kinetic parameters for RDV-resistant viruses showed no dramatic increases in the efficiency of infectious virus production in any mutant virus. But importantly, all mutants rendered RDV ineffective to some extent. Molecular modeling and molecular dynamics simulations of the mutant NSP12 proteins revealed that the tested mutations, excluding F480L, contributed to increased molecular vibrations around the RNA-binding site. Our multidisciplinary approach of molecular virology, mathematical, and molecular modeling discovered mutations involved in RDV resistance and the mechanisms of this drug resistance.

When evaluating virus growth kinetics, titers of different viruses are generally compared at the same time points. However, the total numbers of cells susceptible to viral infection are not constant between wells. The faster a virus propagates and spreads, the faster the number of cells available for virus infection decreases. Thus, comparison of titers at specific time points may not be the best method of evaluating the proliferative capability of viruses with different properties or the effects of drugs against them. Here we combined mathematical models and statistical methods, analyzed intercellular infection dynamics of SARS-CoV-2 in the momently changing cells, and identified the differences in infectious virus production and antiviral effects between WT and mutant viruses (Fig. 3).

Furthermore, molecular simulation of the NSP12 structure clarified the mechanism causing failure of V557L to proliferate and provided new insights into the mechanisms of RDV resistance of the tested mutants (Fig. 4), in contrast to other previous studies that only identified the location of viral mutations. In mutants resistant to RDV, regardless of the location of the mutation, large differences were observed in the RNA binding site when thermodynamic oscillations were compared with WT. Mutations not involved in resistance showed the opposite trend of variation in terms of protein flexibility. We therefore speculate that flexibility in the binding site may be a factor in resistance to RDV.

Three mutations (C799F, E796G, and E802D) markedly diminished the antiviral effect of RDV (Fig. 3C). These mutations did not contribute to efficient virus production but increased the flexibility of the RNA-binding site, which probably enabled these viruses to evade the functional inhibition by RDV binding. The virus with only the C799F mutation was less efficient in virus production than the virus containing all the other mutations identified in this study. Therefore, some mutations, which were observed in genome regions other than NSP12, might restore the efficiency of virus production in the mutant viruses (Fig. 3B). Unlike the mutations found in *in vitro* analyses, D484Y, F480L, and F480L/V557L affected neither virus production nor RDV sensitivity. These results consistent between *in vitro* passages and computer simulations highlight the accuracy and usefulness of our fusion research.

Every mutation observed in *in vitro* passages of SARS-CoV-2 failed to increase the efficiency of infectious virus production. Co-infection competition assay in previous studies revealed that E802D on SARS-CoV-2 or F476L/V553L mutations on MHV decreased fitness (*15, 20*), consistent with our results. Although further characterization is necessary, these results indicate that the observed mutations in this study are unlikely to be persistent in the virus population without RDV or dramatically accelerate the SARS-CoV-2 epidemic. However, because the E802D mutation found initially during *in vitro* passage was actually detected in clinical patients (*19, 20*), it is quite possible that the analyzed mutations may be reported following the sustained administration of RDV, and thus continuous viral sequence analyses from SARS-CoV-2 patients treated with RDV is vital. Currently, we cannot definitively state that the repeated administration of RDV will be problematic, but additional compounds with higher affinity for the RdRp complex than RDV are likely to become desirable.

The SARS-CoV-2 pandemic is not coming to an end, rather, new virus strains have sequentially emerged. Deletions and mutations that can facilitate higher transmissibility or antibody evasion have been highlighted in the Omicron variant (*29–33*). It has been revealed that two doses of mRNA vaccine are insufficient to provide immunity against the Omicron variant (*34–36*), and new vaccines that will provide more robust immunity are urgently needed. In addition, new antivirals have been developed and are expected to be approved for clinical use (*37–39*). While development of new vaccines and antivirals will continue, it remains important to evaluate their safety, and to pay attention to resistant mutations. Our state-of-the-art study, in which the drug sensitivities of multiple mutant viruses were simultaneously determined, will accelerate the development of new measures, evaluation of drug resistance and deepen our understanding of the driving forces for mutation of SARS-CoV-2.

## Materials and methods

### Cells and viruses

HEK293-C34 cells were previously established and a different clone than HEK293-3P6C33 cell, both of which were IFNAR1 deficient, with expression of human ACE2 and TMPRSS2 induced by doxycycline hydrochloride (*21*). The HEK293-C34 cells were maintained in Dulbecco’s Modified Eagle Medium (DMEM) (Nacalai Tesque) containing 10% fetal bovine serum (FBS) (Sigma) and blasticidin (10 μg/ml) (Invivogen). The exogenous expression of ACE2 and TMPRSS2 in the HEK293-C34 cells was induced by addition of doxycycline hydrochloride (1 μg/ml) (Sigma). TMPRSS2-expressing Vero E6 (VeroE6/TMPRSS2) cells were purchased from the Japanese Collection of Research Bioresources Cell Bank (JCRB1819) and maintained in DMEM containing 10% FBS and G418 (Nacalai Tesque). Both HEK293-C34 cells and VeroE6/TMPRSS2 cells were cultured at 37°C in 5% CO2.

SARS-CoV-2 strain JPN/TY/WK-521 was kindly provided by Dr. Masayuki Shimojima at the National Institute of Infectious Diseases. All experiments involving SARS-CoV-2 were performed in biosafety level 3 laboratories, following the standard biosafety protocols approved by the Research Institute for Microbial Diseases at Osaka University.

### Chemical inhibitors and antibodies

RDV (GS-5734) was purchased from Cayman Chemical, dissolved in dimethyl sulfoxide (DMSO) and stored at 50 mM at −30°C. To detect SARS-CoV-2-infected cells, mouse monoclonal antibody against SARS-CoV-2 NP (Clone# S2N4-1242) was kindly provided by Bio Matrix research. Alexa Fluor 488-conjugated anti-mouse antibodies were purchased from Life Technologies.

### Serial passages of SARS-CoV-2

SARS-CoV-2 strain JPN/TY/WK-521 was serially passaged in HEK293-C34 cells 10 times in the presence of RDV (Fig. 1). HEK293-C34 cells were prepared with DMEM containing 10% FBS, blasticidin, and 1 μg/ml doxycycline hydrochloride in six-well plates. One day later, the cells were infected with SARS-CoV-2 at a multiplicity of infection of 0.01 for 1 hour to allow virus attachment. Culture supernatants were then replaced with DMEM containing 2% FBS, blasticidin, 1 μg/ml doxycycline hydrochloride, and RDV. The virus-infected cells were incubated and the supernatants were collected when CPE was observed throughout the wells. The collected supernatants were centrifuged at 1,500 ×g for 5 min to remove cells and debris, and 10 μM of the supernatants were passaged. The final concentration of RDV was gradually increased from 0.01 μM in P1 to 4.0 μM in P10.

### Validation of the virus sequence

The virus sequences were confirmed by Sanger sequencing. Total RNA was extracted from the supernatants of SARS-CoV-2-infected cells by using a PureLink RNA mini kit (Invitrogen) and subjected to cDNA synthesis using a PrimeScript RT reagent kit (Perfect Real Time) (TaKaRa Bio) and random hexamer primers. A total of nine DNA fragments, covering the full-length SARS-CoV- 2 genome were amplified by PCR using a PrimeSTAR GXL DNA polymerase (TaKaRa Bio), the synthesized cDNA and specific primer sets from CoV-2-G1-Fw to CoV-2-G10-Rv designed previously (*21*). The amplified PCR fragments were purified using a gel/PCR DNA isolation system (Viogene) and sequenced in both directions using the ABI PRISM 3130 genetic analyzer (Applied Biosystems) with specific primers for SARS-CoV-2.

### Rescue of the presumed RDV-resistant viruses

All the HiBiT-carrying SARS-CoV-2 with RDV-resistant mutations were rescued by the CPER method, which was established in our previous study (*21*). Briefly, nine cDNA fragments, covering the entire genome of SARS-CoV-2 were prepared by PCR using PrimeSTAR GXL DNA polymerase and SARS-CoV-2 viral gene fragment-encoding plasmids. In addition, an untranslated region (UTR) linker fragment encoding the 3′ 43 nucleotides (nt) of SARS-CoV-2, hepatitis delta virus ribozyme (HDVr), bovine growth hormone (BGH) poly(A) signal, cytomegalovirus (CMV) promoter, and the 5′ 25 nt of SARS-CoV-2, was amplified by PCR. A HiBiT luciferase gene (VSGWRLFKKIS) and a linker sequence (GSSG) were introduced into the N terminus of the ORF6 gene of SARS-CoV-2 by site-directed mutagenesis of the viral genome fragment-cloning plasmid and the plasmid was used as a template to amplify a cDNA fragment. Presumed RDV-resistant mutations were introduced into the SARS-CoV-2 cDNA fragments by overlap PCR using specific overlapping primer sets. The nine SARS-CoV-2 cDNA fragments and the UTR linker fragment (0.1 pmol each) were mixed together and subjected to CPER. The CPER products were then directly transfected into HEK293-C34 cells using Trans IT LT-1 (Mirus). At 6 hours post-transfection, the culture media were changed to DMEM containing 2% FBS, blasticidin, and doxycycline hydrochloride (1 mg/ml). When CPE was observed throughout the wells (usually around 7 days post-transfection), the culture supernatants were collected and centrifuged at 1,500 ×g for 5 min to remove cells and debris. The culture supernatants were then passaged once using VeroE6/TMPRSS2. The virus sequences were confirmed in the passaged virus solutions by Sanger sequencing and thereafter the virus stocks were stored at −80°C until use.

### Virus titration

Infectious titers in culture supernatants were determined by 50% tissue culture infective doses (TCID50). VeroE6/TMPRSS2 cells were prepared in 96-well plates and infected with SARS-CoV-2 after ten-fold serial dilution with DMEM containing 2% FBS. Virus titers were determined at 72 hpi.

### RDV susceptibility analysis using the HiBiT system

HEK293-C34 cells were seeded in 48-well plates in DMEM with 10% FBS, blasticidin and 1 μg/ml doxycycline hydrochloride. One day later, HiBiT-carrying SARS-CoV-2 was allowed to attach for 1 hour. Culture media were then replaced with new media containing 2% FBS, blasticidin, 1 μg/ml doxycycline hydrochloride, and RDV (0–1.0-μM final concentration). At 48 hpi, luciferase activity was measured using a Nano-Glo HiBiT lytic assay system (Promega), following the manufacturer’s protocols. Briefly, Nano-Glo substrate including LgBiT protein was added to the virus-infected cell lysates after all culture supernatants were removed. Luciferase activities were measured using a luminometer and normalized to luminescence without RDV treatment (0-μM final concentration). The EC50 was calculated using the drc package (v3.0-1; R Project for Statistical Computing).

### Time course analyses of infectious virus production

HEK293-C34 cells were prepared in 96-well plates in media containing 1 μg/ml doxycycline hydrochloride. Cells were infected with HiBiT-carrying viruses at MOI=0.01 for 1 hour. Culture media were changed to fresh media containing 2% FBS, blasticidin, and 1 μg/ml doxycycline hydrochloride, with or without 0.05 μM RDV (final concentration). At 12, 24, 48, 72, and 96 hpi, culture supernatants of the virus-infected cells were collected and infectious titers in the supernatants (TCID50/ml) were determined by virus titration.

### Time course analyses of infection rates in cells

After removal of the culture supernatants at the indicated time points in the time course analyses of infectious virus production, the cells were fixed with 4% paraformaldehyde (Nacalai Tesque). The fixed cells were permeabilized with 0.2% Triton X-100 (Nacalai Tesque) in PBS for 20 min, blocked with 1% bovine serum albumin fraction V (Sigma) in PBS, and then reacted with anti-SARS-CoV-2 NP antibody in PBS for 1 hour at room temperature. After washing with PBS three times, the cells were incubated with a 1:1,000 dilution of goat anti-mouse IgG Alexa Fluor 488- conjugated secondary antibody (Thermo Fisher Scientific) in PBS for 1 hour at room temperature. The cells were then incubated with DAPI (Thermo Fisher Scientific) (1:2,000 dilution) for 10 min. Immunopositive signals were confirmed under a FluoView FV1000 confocal laser scanning microscope (Olympus), with appropriate barrier and excitation filters. Quantitative imaging data were obtained using a CellVoyager CQ1 benchtop high-content analysis system (Yokogawa Electric Corporation) and analyzed with CellPathfinder high content analysis software (Yokogawa Electric Corporation). The number of SARS-CoV-2-infected cells stained by anti-SARS-CoV-2 NP antibody and the number of cell nuclei stained by DAPI were counted. The infection rates were then calculated by dividing the number of SARS-CoV-2-positive cells by the total number of cell nuclei.

### Growth of HEK293-C34 cells

HEK293-C34 cells were seeded in 48-well plates with DMEM containing 10% FBS and blasticidin. At 24 hours post-seeding, media were replaced with DMEM containing 10% FBS, blasticidin, and 1 μg/ml doxycycline. All the cells were collected, and the cell numbers were counted every 12 hours for 48 hours post-medium change.

### Degradation rate of HiBiT-carrying SARS-CoV-2

HiBiT-carrying SARS-CoV-2 were incubated at 37°C with 5% CO2 for 48 hours. Every 12 hours, the virus solutions were collected and subjected to virus titration to quantify infectious virus.

### Quantification of cell growth and virus decay kinetics

To estimate the growth kinetics of target cells, we used the following mathematical model:

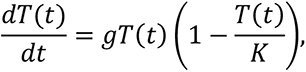

where the variable *T*(*t*) represents the number of uninfected target cells (cells/ml) at time *t*, and the parameters *g* and *K* indicate the growth rate and the carrying capacity of the target cells (cells/ml), respectively. Using the non-linear least square method, we fitted the model to the time-course growth data of cells (see **Growth of HEK293-C34 cells** and **Fig. S2A**) and estimated *g* and *K*.

Furthermore, we estimated the clearance rate of infectious viruses, *c*, by a simple exponential decay model:

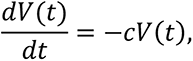

where *V*(*t*) represents the amount of infectious virus (TCID50/ml) in the culture medium at time *t*. Linear regressions yield *c* from the time-course degradation data of infectious viruses (see **Degradation rate of HiBiT-carrying SARS-CoV-2** and **Fig. S2B**). The estimated parameter values are summarized in **Table 2**.

### Mathematical model for SARS-CoV-2 infection

We employed the following mathematical model for SARS-CoV-2 infection in cell culture considering the antiviral efficacy of RDV:

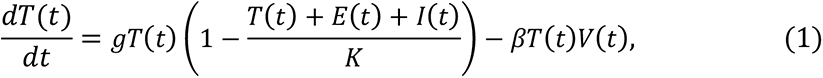

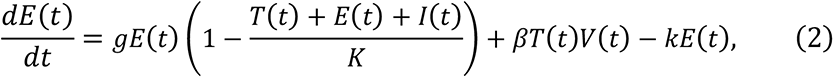

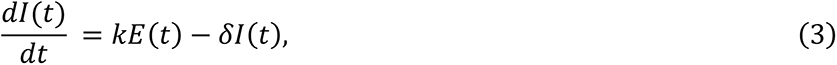

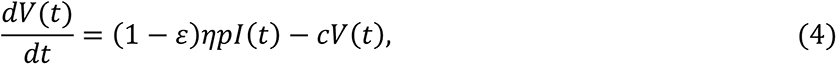

where *T*(*t*), *E*(*t*), and *I*(*t*) are the numbers of uninfected target cells, eclipse phase cells, and virus-producing cells (cells/ml) at time *t*, respectively, and *V*(*t*) is the amount of infectious virus (TCID50/ml) at time *t*. The uninfected target and eclipse phase cells divide in logistic manner at rate *g* and carrying capacity *K*. The target cells are infected by viruses at rate *β*, and the virus-infected cells stay in the eclipse phase during the period 1/*k*. After this, they become virus-producing cells. The progeny viruses are produced by the virus-producing cells at rate 𝑝. The parameters 𝛿 and *c* indicate the death rate of infected cells and the clearance rate of viruses, respectively. The inhibition rate of virus production by RDV is assumed to be 𝜀. The fold-change of virus production rates of RDV-resistant viruses compared with WT virus are 𝜂 (i.e., 𝜂 = 1 for WT virus).

### Data fitting and parameter estimation

The parameters *g*, *K*, and *c* were independently estimated and fixed. A statistical model adopted in Bayesian inference assumed that measurement error followed a normal distribution with mean zero and constant variance (error variance). A gamma distribution was used as a prior distribution, and it inferred a distribution of error variance. As an output of MCMC computations, the posterior predictive parameter distribution represented parameter variability, and it inferred distributions of model parameters and initial values of variables. The estimated parameters and initial values are listed in **Table 2** and **Table 3**.

### Theoretical predictions and analyses of the effects of amino acid mutations

The WT NSP12 structure used as a reference was created based on the crystal structure (PDB ID: 6XEZ) (*40*). From this crystal structure, only the NSP12 protein and 28 bases of the binding RNA (binding site) were extracted. The target mutant structures were constructed by substituting amino acids in the above WT structure. Based on the crystal structure (PDB ID: 7BV2) (*41*), the terminal nucleotides of each predicted structure were replaced with RDV. To neutralize the charge of these complex structures, counter ions were placed, and sufficient water molecules were placed around them. Each structure was stabilized by the energy minimization method and used as the initial structure for molecular dynamics simulations. The composite structure was thermally stabilized by raising the temperature from 0 K to 310 K (*in vivo* temperature, approximately 36.85°C) over 500,000 steps with Δt = 0.2 fs. The structural changes during this temperature increase process were structurally sampled for each complex at every 1000 steps. These sampling structures were superimposed, and the RMSD for each protein was calculated. In addition, to observe the extent to which the structural properties differed between WT and mutants, we superimposed them in various substructures and calculated and compared the RMSD differences. These energy minimizations and molecular dynamics simulations were performed with the AMBER18 program package (*42*). The “AMBER99 (*43*)”, “GAFF (*44*)” and “TIP3P (*45*)” force fields for the “proteins and nucleic acids”, “RDV”, and “water molecules” were employed, respectively.

### Statistical analysis

Results are indicated as the means ± standard deviations or standard errors. Statistical significances were determined by the one-way ANOVA with Dunnett’s test or the Kruskal–Wallis test with the two stage linear step-up procedure of Benjamini, Kreiger, and Yekutieli, which was performed using GraphPad Prism (Software ver. 9.2.0). Significantly different values are indicated by asterisks (\**p*<0.05 or \*\*\**p*<0.001).

## Acknowledgments

We thank M. Tomiyama for her secretarial work, M. Ishibashi and K. Toyoda for their technical assistance. We also thank Dr. Dr. Masayuki Shimojima at NIID for providing SARS-CoV-2 strain JPN/TY/WK-521 and Bio Matrix Research for providing anti-SARS-CoV-2 NP monoclonal antibodies. This study was supported in part by a Grant-in-Aid for JSPS Scientific Research (KAKENHI) B 21H02736 (to T.F.), 18KT0018 (to S. Iwami), 18H01139 (to S. Iwami), 16H04845 (to S. Iwami), 20H04281 (to T.S.), Scientific Research in Innovative Areas 20H05042 (to S. Iwami), 19H04839 (to S. Iwami), 18H05103 (to S. Iwami), 20H04841 (to T.S.); AMED CREST 19gm1310002 (to S. Iwami); AMED Japan Program for Infectious Diseases Research and Infrastructure, 20wm0325007h0001 (to S. Iwami), 20wm0325004s0201 (to S. Iwami), 20wm0325012s0301 (to S. Iwami), 20wm0325015s0301 (to S. Iwami); AMED Research Program on HIV/AIDS 19fk0410023s0101 (to S. Iwami); AMED Research Program on Emerging and Re-emerging Infectious Diseases 20fk0108401 (to T.F.), 20fk010847 (to T.F.) 21fk0108617 (to T.F.), 20fk0108451 (to T.F.), 20wm0325007s0201 (to S.H.), 19fk0108050h0003 (to S. Iwami), 19fk0108156h0001 (to S. Iwami), 20fk0108140s0801 (to S. Iwami) and 20fk0108413s0301 (to S. Iwami); AMED Program for Basic and Clinical Research on Hepatitis 19fk0210036h0502 (to S. Iwami); AMED Program on the Innovative Development and the Application of New Drugs for Hepatitis B 19fk0310114h0103 (to S. Iwami); JST MIRAI (to S. Iwami); Moonshot R&D Grant Number JPMJMS2021 (to S. Iwami) and JPMJMS2025 (to S. Iwami); Mitsui Life Social Welfare Foundation (to S. Iwami); Shin-Nihon of Advanced Medical Research (to S. Iwami); Suzuken Memorial Foundation (to S. Iwami); Life Science Foundation of Japan (to S. Iwami); SECOM Science and Technology Foundation (to S. Iwami); The Japan Prize Foundation (to S. Iwami); Daiwa Securities Health Foundation (to S. Iwami). We thank Gillian Campbell, PhD, from Edanz (https://www.jp.edanz.com/ac), for editing a draft of this manuscript.

---

**Fig. S1.**
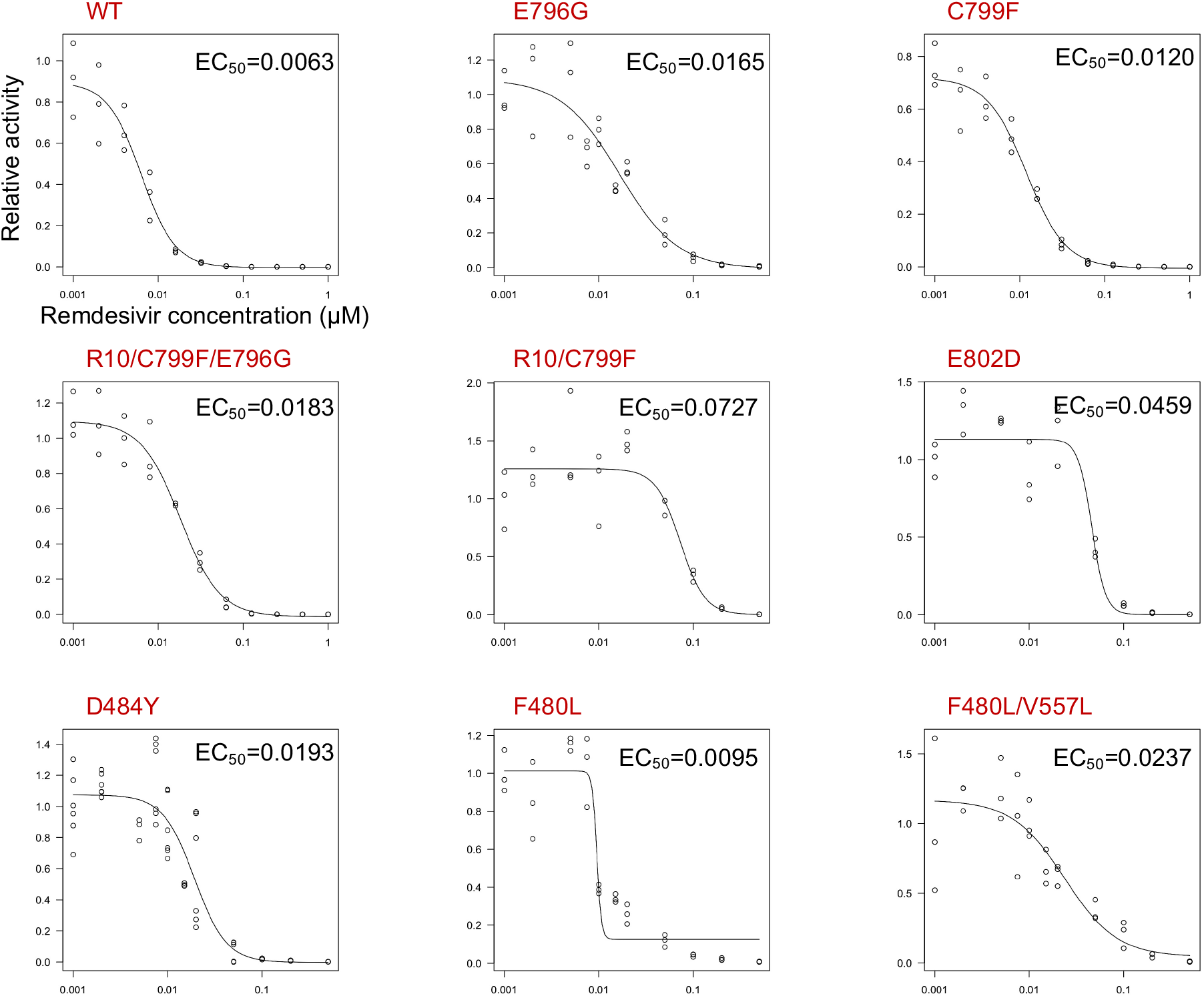
RDV susceptibility of the mutants. Cells were infected with HiBiT-carrying SARS-CoV- 2 viruses in the presence or absence of RDV (0–1.0 μM final concentration) for 48 hours.Luciferase activities were measured and normalized to no RDV treatment. EC50 was calculated using the drc package (v3.0-1).

**Fig. S2.**
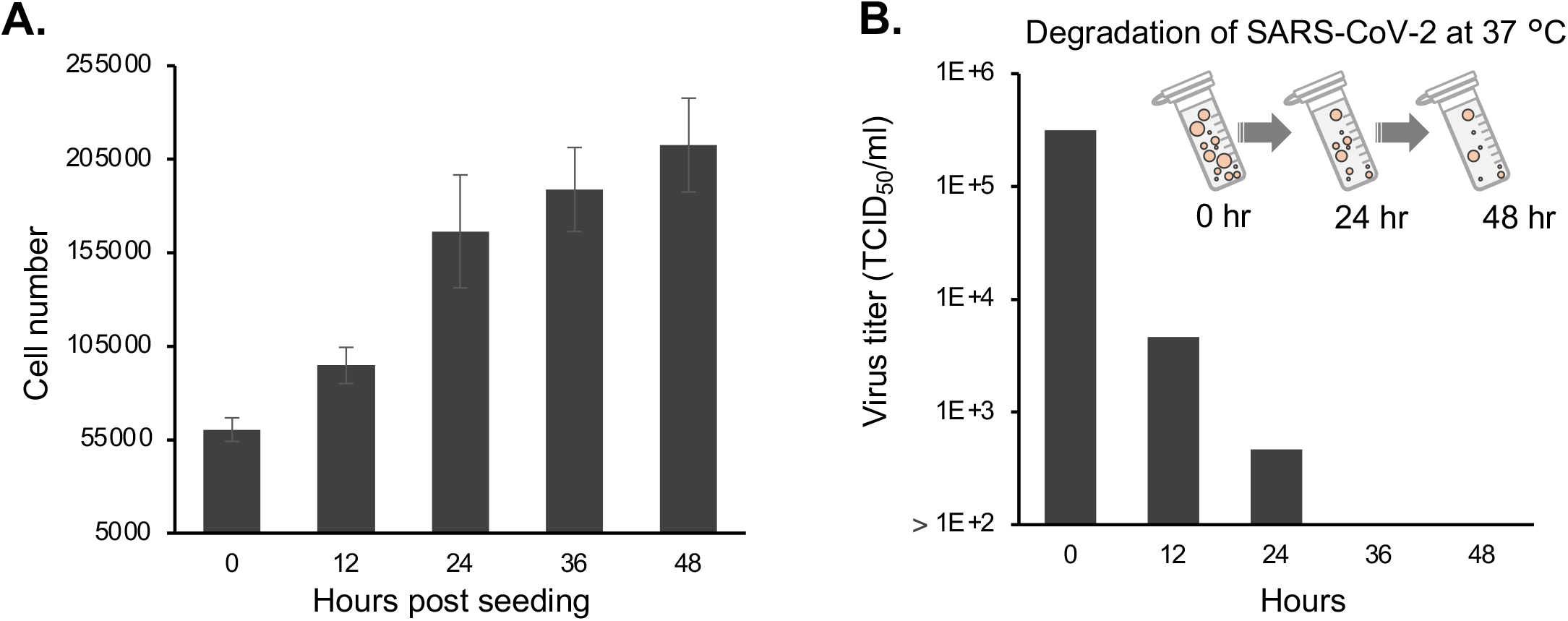
Biological characterization of HEK293-C34 cells and SARS-CoV-2. **(A)** Growth kinetics of HEK293-C34 cells. HEK293-C34 cells were counted for 48 hours after seeding. **(B)** Degradation rate of SARS-CoV-2. Virus titers were determined every 12 hours during incubation at 37°C.

**Fig. S3.**
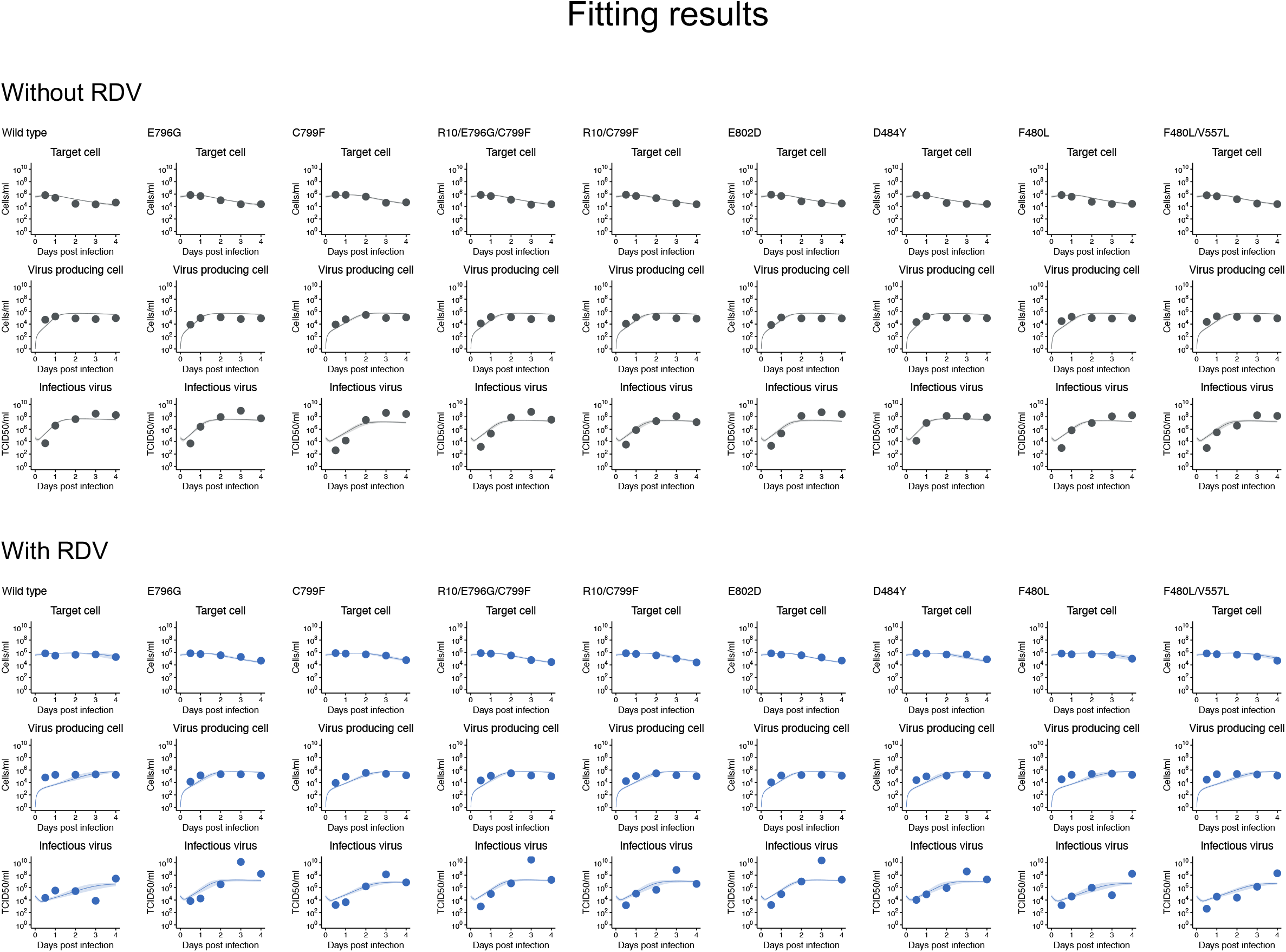
SARS-CoV-2 infection dynamics without and with RDV. Solid curves and shadowed regions correspond to the best-fit solution and 95% posterior intervals, respectively, of equations (1–4) for the time-course dataset (black and blue dots). Top and bottom panels correspond to experiments without and with RDV treatment, respectively. All data for each strain were fitted simultaneously.

**Fig. S4.**
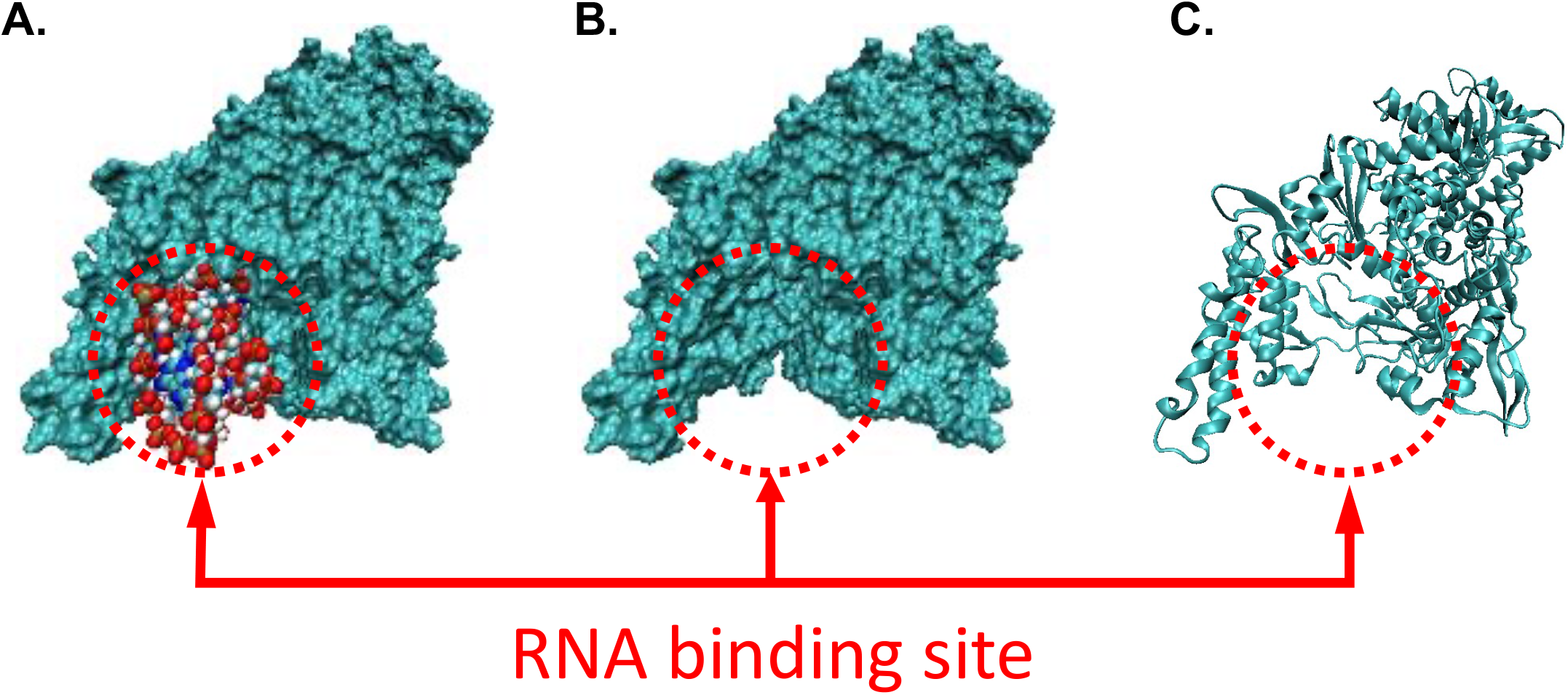
RNA binding sites in NSP12 protein. **(A)** Surface notation for NSP12 protein and van der Waals notation for bound RNA**. (B and C)** The RNA binding space is exposed by using only the protein surface notation and cartoon notation.

